## Supplemental Figures for "Training-induced bioenergetic improvement in human skeletal muscle is associated with non-stoichiometric changes in the mitochondrial proteome without reorganization of respiratory chain content"

### SUPPLEMENTARY FIGURES

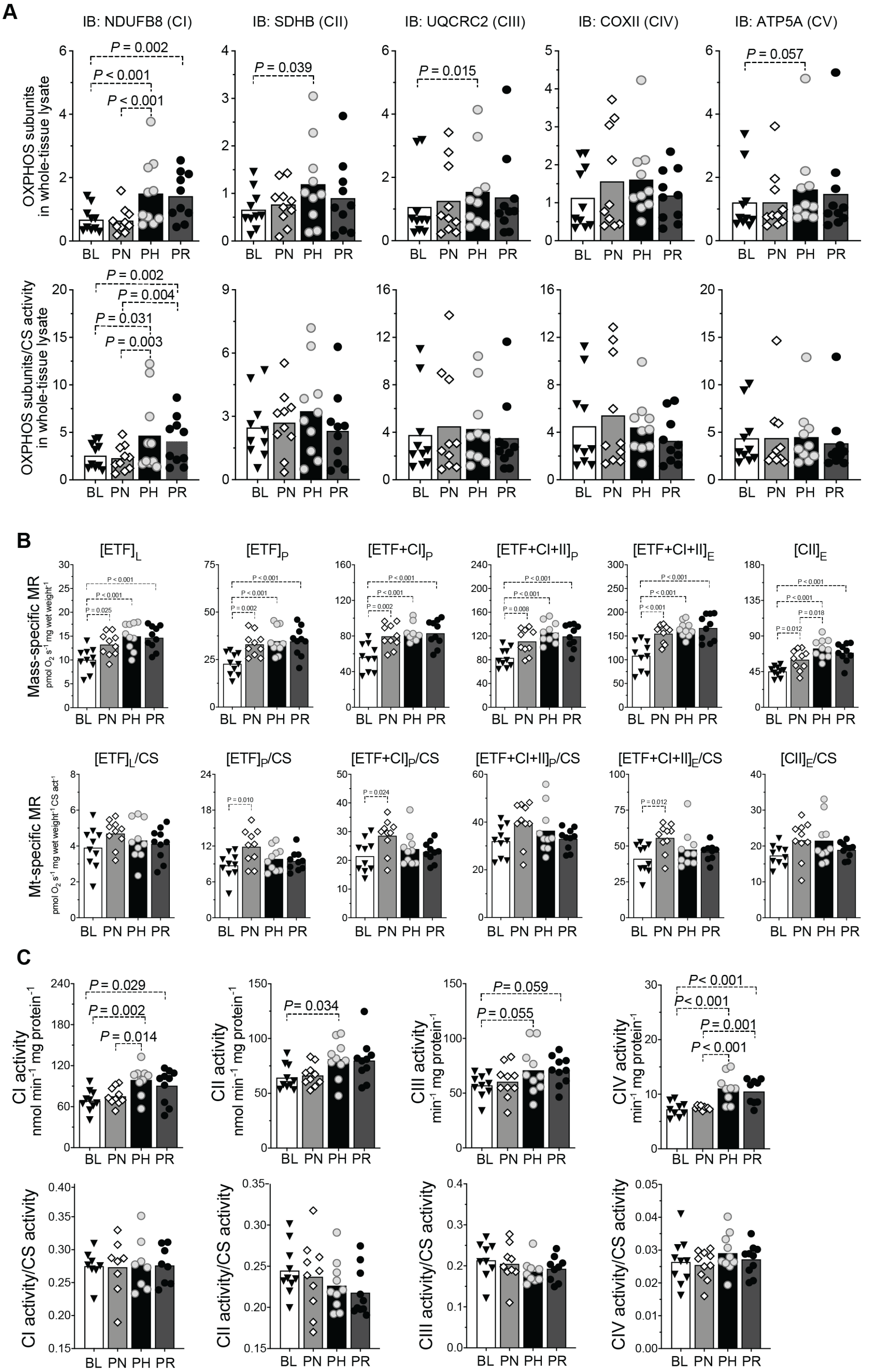


**Figure S1**.

(A) Top panels: protein content of selected subunits of oxidative phosphorylation (OXPHOS) complexes by immunoblotting in whole-tissue (vastus lateralis) homogenates; lower panels: values from top panels normalised by citrate synthase (CS) activity (obtained in Figure 1D). (B) Top panels: mass-specific mitochondrial respiration (MR) in permeabilized human vastus lateralis muscle fibers measured with the following substrate-uncoupler-inhibitor titration (SUIT) protocol: $[\mathrm{ETF}]_L$, leak respiration state (L) in absence of adenylates and electron input through ETF; $[\mathrm{ETF}]_P$, maximal OXPHOS state (P) with electron input through ETF; $[\mathrm{ETF+CI}]_P$, P with convergent electron input through ETF + CI; $[\mathrm{ETF+C+II}]_P$, P with convergent electron input through ETF + CI + CII; $[\mathrm{ETF+C+II}]_E$, maximal electron transport chain capacity (E) with convergent electron input through ETF + CI + CII; $[\mathrm{CII}]_E$, E with electron input through CII. Lower panels: mitochondrial (mt)-specific MR obtained by normalizing values of mass-specific MR by CS activity (obtained in Figure 1D). (C) Top panels: enzymatic activity of electron transport chain (ETC) complexes in whole-tissue (vastus lateralis) homogenates; lower panels: values from top panels normalised by CS activity (obtained in Figure 1D).

BL: baseline; PN: post-NVT; PH: post-HVT; PR: post-RVT; CI-V: complex I to V; IB: immunoblotting; ▼,◊,○, and ● represent individual values; bars represent mean values; n = 10 for all analyses. All datasets analyzed by repeated measures one-way ANOVA followed by Tukey’s post hoc testing, except for $[\mathrm{ETF+C+II}]_P$/CS, which was analyzed by Friedman test followed by Dunn’s post hoc testing, as not normally distributed. Significance: $P < 0.05$.

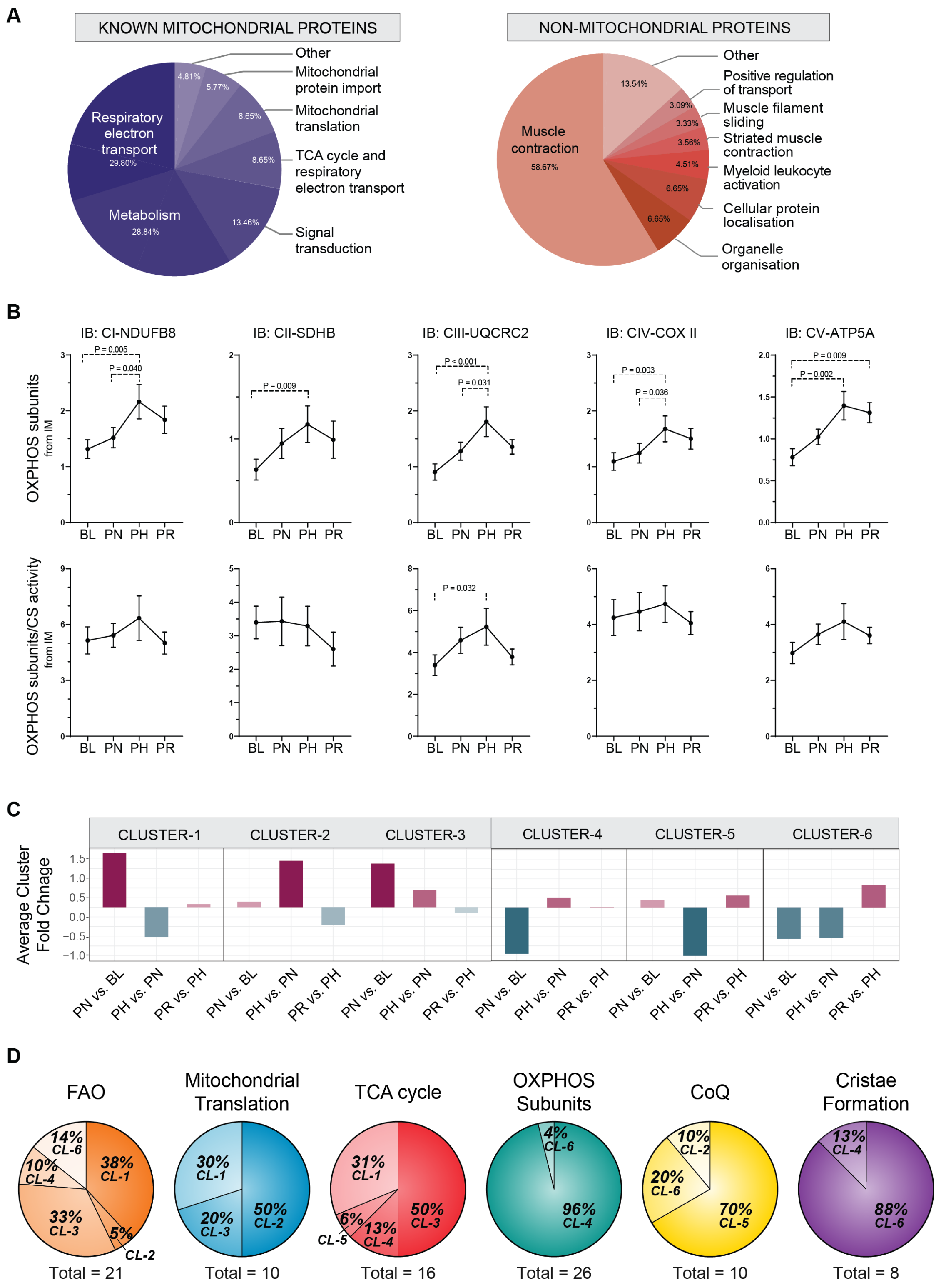


**Figure S2.**

(A) Pie charts showing the relative enrichment terms, as determined by *Reactome*, of "Known Mitochondrial" and "non-mitochondrial" proteins, identified by the Integrated

Mitochondrial Protein Index (IMPI) database (Smith and Robinson, 2019) from isolated mitochondria (IM) fractions.

(B) Top panels: training-induced changes in protein content of selected subunits of oxidative phosphorylation (OXPHOS) complexes by SDS-PAGE in IM fractions from vastus lateralis muscle biopsies; lower panels: values from top panels normalized by CS activity (obtained in Figure 1D).

(C) Profile plots of the scaled expression mean of proteins within the six clusters as determined in Figure 2E.

(D) Venn diagram representation of the protein distribution from each of the six main protein functional classes (as determined in Table S7) within the six clusters as determined in Figure 2E.

BL: baseline; PN: post-NVT; PH: post-HVT; PR: post-RVT; IB: immunoblotting; n = 10 for all analyses; all datasets analyzed by repeated measures one-way ANOVA followed by Tukey's post hoc testing. Significance: $P < 0.05$.

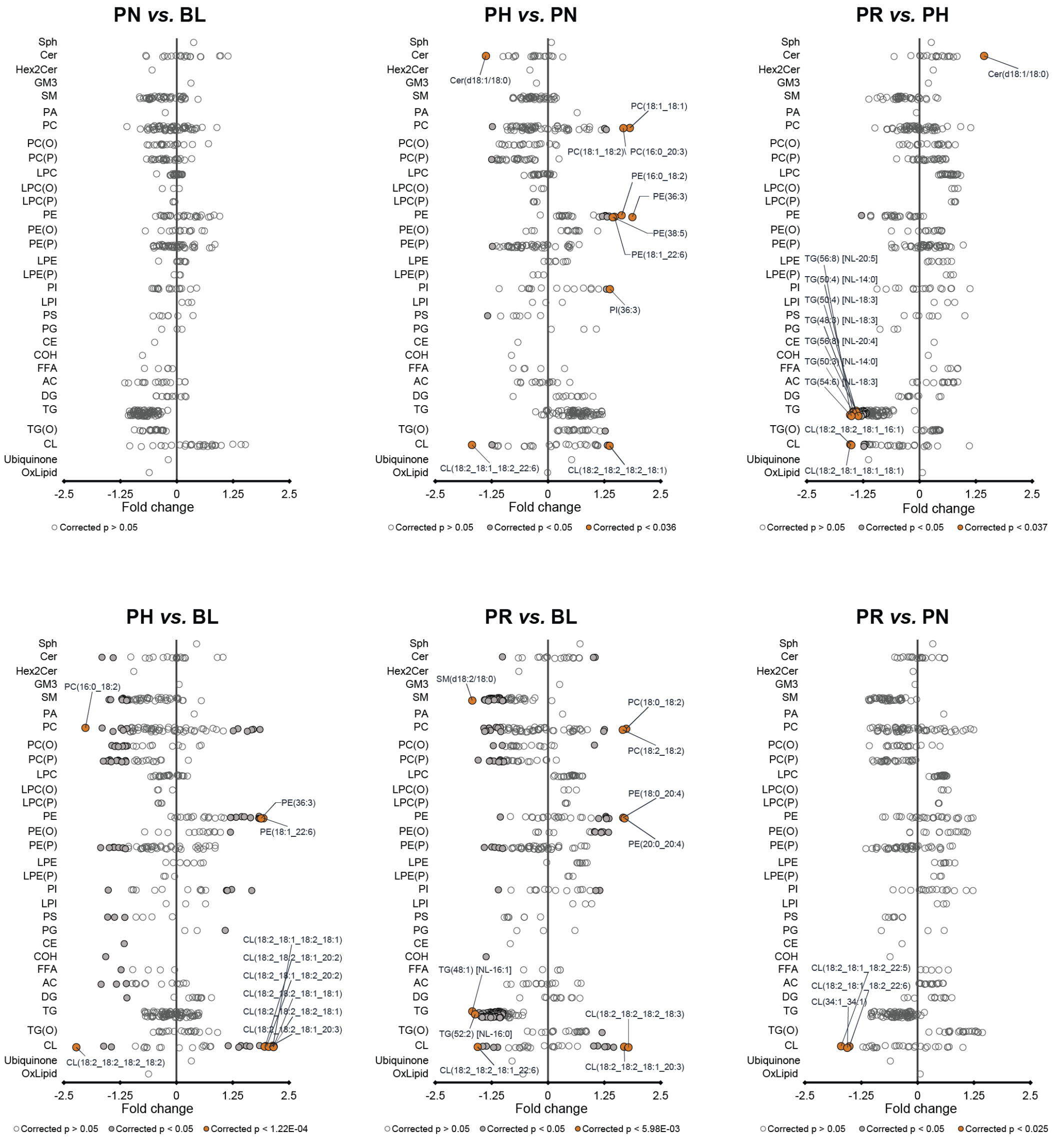


**Figure S3**.

Forest plots displaying training-induced changes in individual lipid species between pairs of time points obtained from lipidomics analysis of isolated mitochondria (IM) fractions from human vastus lateralis muscle biopsies; n = 10 for all analyses; all data from differential expression tests taken from *Limma*; open circles show non-significant species, gray circles show species with $P < 0.05$ after correction for multiple comparisons (Benjamini-Hochberg). The top 10 species after correction for multiple comparisons are highlighted with orange circles. BL: baseline; PN: post-NVT; PH: post-HVT; PR: post-RVT.
